# Red yeast rice-derived MKA ameliorates cardiac hypertrophy in hypertensive rats by inhibiting ERK1/2/c-Fos pathway

**DOI:** 10.64898/2026.03.10.710945

**Authors:** Rubin Tan, Dongqi Yang, Kuntao Liu, Jia Liu, Na Li, Manqi Sun, Xiaoqiu Tan, Qinghua Hu, Chunxiang Zhang

## Abstract

**Background:** Cardiac hypertrophy is a key pathological process in hypertensive heart failure, yet current antihypertensive therapies do not directly target it. Red yeast rice (RYR), rich in monacolin K β-hydroxy acid (MKA), is known for lipid-lowering effects, but its potential to ameliorate cardiac hypertrophy is unreported.

**Purpose:** To investigate the effects of RYR-derived MKA on cardiac hypertrophy in spontaneously hypertensive rats (SHR) and elucidate its molecular mechanisms.

**Methods:** Spontaneously hypertensive rats (SHR) were treated with 0.6% red yeast rice for 8 weeks to assess its effects on blood pressure, cardiac function (echocardiography), cardiac hypertrophy and fibrosis (histopathology), and multi-organ toxicity (histopathology). A multigenerational study was conducted to evaluate protective effects in offspring. Network pharmacology and transcriptomic analysis were integrated to predict molecular targets, which were subsequently validated by molecular docking and experiments.

**Results:** Eight-week RYR treatment significantly reduced blood pressure, inhibited cardiac hypertrophy and fibrosis, and improved cardiac function without gender differences. No pulmonary, hepatic, or renal toxicity was observed. Offspring from treated parents exhibited further reduced hypertrophy upon continued treatment. Mechanistically, MKA bound ERK1/2 with high affinity, inhibiting its phosphorylation and downstream c-Fos expression, thereby downregulating hypertrophy markers.

**Conclusion:** Red yeast rice improves hypertensive cardiac hypertrophy via MKA-mediated inhibition of the ERK1/2/c-Fos pathway. Its multi-organ safety and transgenerational effects offer a novel dual-therapy strategy for hypertension and cardiac hypertrophy.

**Graphic abstract:** 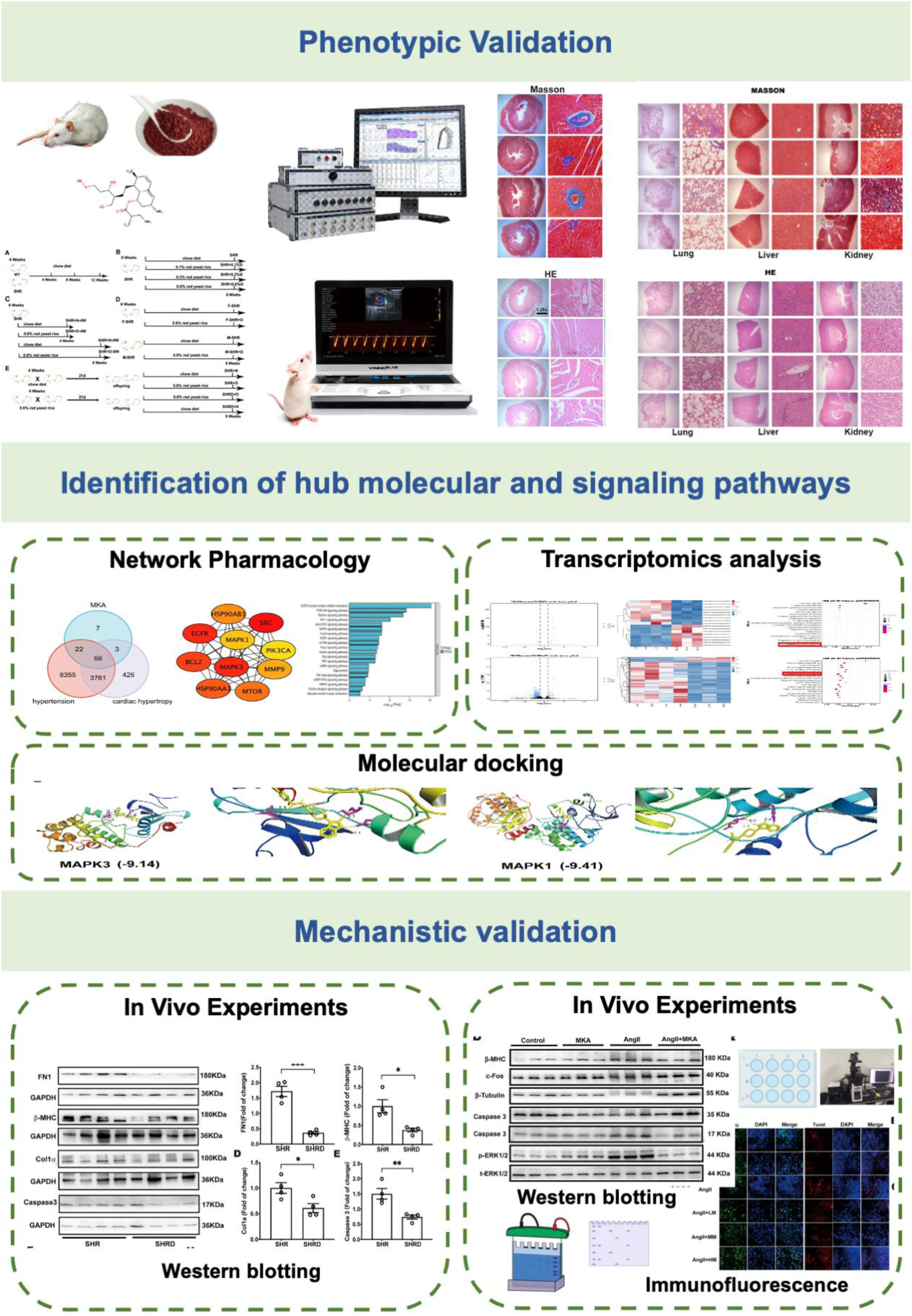

## Introduction

Hypertension is a prevalent chronic condition primarily characterized by elevated systemic arterial blood pressure (Te Riet et al., 2015). Its global burden is estimated at 874 million patients, with approximately 9.4 million cardiovascular disease deaths annually attributable to the condition (Forouzanfar et al., 2017). One clinical consequence of hypertension is left ventricular hypertrophy and myocardial remodeling, which can precipitate heart failure (González et al., 2018; Yildiz et al., 2020). Current guidelines recommend antihypertensive medication for treating cardiac hypertrophy in hypertensive patients (Rabi et al., 2020). Currently available antihypertensive agents include agents acting on the renin-angiotensin system (ACEIs), β-blockers, calcium channel blockers (CCBs), and diuretics. A randomized controlled trial by Leache et al. (2021) found no benefit in cardiovascular event rates or mortality among hypertensive participants with left ventricular hypertrophy when these antihypertensive drugs were added. Concurrently, in clinical practice, while these commonly prescribed oral antihypertensive agents alleviate symptoms, they frequently induce adverse reactions: ACEIs may cause irritating dry cough and angioedema (AlQudah et al., 2020); β-blockers may exacerbate atrioventricular block and offer no benefit to asthma patients (Argulian et al., 2019); CCBs may induce headaches and ankle oedema in certain patients; high-dose diuretics affect water, electrolyte, and metabolic balance (Blowey, 2016; Burnier et al., 2019). Given these limitations and the absence of interventions specifically targeting myocardial hypertrophy, the necessity for exploring novel therapeutic approaches is highlighted.

In recent years, an increasing body of research has demonstrated that traditional Chinese medicine can protect vascular endothelial function while lowering blood pressure, thereby offering unique advantages in preventing and treating complications of hypertension and improving patients’ quality of life. *Red yeast rice* has been widely employed in traditional Chinese medicine, health supplements, and food products in China. For instance, a randomized controlled trial in myocardial infarction patients demonstrated that *Xuezhikang* capsules containing *red yeast rice* significantly reduced the risk of major coronary events by 45% (from 10.4% in the placebo group to 5.7%), decreased cardiovascular mortality by 30%, and reduced all-cause mortality by 33%. while improving total cholesterol, low-density lipoprotein cholesterol (LDL-C), and triglyceride levels with good tolerability (Lu et al., 2008). Long-term administration of Lipid-Kang reduces coronary heart disease events and improves lipid profiles in Chinese myocardial infarction patients, demonstrating safety and efficacy (Lu et al., 2008). Chromatographic analysis further identifies monacolin K β-hydroxy acid (MKA) as the primary active component of *red yeast rice* (Higa et al., 2021; Huang et al., 2006). Daily intake of low-dose *red yeast rice* containing 2mg MKA significantly reduced LDL-C, total cholesterol, apolipoprotein B, and blood pressure levels in patients with mild dyslipidemia, without muscle or hepatic/renal toxicity, indicating its potential for cardiovascular risk management (Minamizuka et al., 2021). Another study also found that A 60-day treatment with low-dose MKA dietary supplements significantly reduced LDL-C and total cholesterol by 15.6% and 15.3%, respectively, in patients with moderate-to-severe dyslipidemia, with non-HDL cholesterol decreasing by 35.4%, without inducing serious adverse events (Benjian et al., 2022). However, no reports exist on whether *red yeast rice* directly targets cardiomyocytes to improve hypertensive cardiac hypertrophy.

This study utilizes animal experiments to clarify red yeast rice’s efficacy in reducing hypertension and improving hypertensive cardiac hypertrophy, conducting multidimensional evaluations of its toxic side effects (male and female individuals, parental and offspring generations, treatment duration, multiple organs). Network pharmacology combined with transcriptomics analysis identifies MKA-derived targets and molecular mechanisms from red yeast rice, with animal and cellular experiments validating MKA’s molecular mechanisms for improving hypertensive myocardial hypertrophy.

## 2. Materials and Methods

### 2.1 Animal Models

Wista rats (control) and spontaneously hypertensive rats (SHR) were purchased from ^©^Charles River. The animal research protocol was reviewed and approved by the XZHMU Animal Care and Use Committee (Ethical Approval No. 202208S058), with all procedures conducted in accordance with the Guidelines for the Care and Use of Laboratory Animals.

### 2.2 Experimental Design

*Red yeast rice*, predominated by the MKA in form of micropowder, purchased from Tongjuntan®, Hangzhou, Zhejiang, China. The total MKA content is around 3.5 mg in per gram red yeast rice (Higa et al., 2021; Huang et al., 2006). Five animal experiments were designed for in vivo experiments (***Figure. S1,*** n=5 for each group).

### 2.3 Cardiac echocardiography

SHR were administered 3% pentobarbital sodium (30 mg/kg) via intraperitoneal injection. Apply coupling agent to the depilated thoracic area. Assessment of cardiac function through transthoracic echocardiography using VINNO 6LAB ultrasound equipment (VINNO Technology Co., Ltd, China). Use M-type long axis view to evaluate left ventricular end diastolic volume (LVEDV), left ventricular end systolic volume (LVESV), left ventricular ejection fraction (EF) and left ventricular fractional shortening (FS). This process adopts a double-blind experiment.

### 2.4 Hemodynamic analysis

Blood pressure was measured through Left common carotid artery catheterization, and the amplitude and frequency of the waveform were observed to confirm normality.

### 2.5 Harvest of tissue samples

After hemodynamic analysis, rats were then sacrificed. The hearts, lungs, livers and kidneys are harvested to do the next experiments.

### 2.6 Hematoxylin & eosin and Masson staining

Hematoxylin and eosin (H&E) and Masson staining were performed as our previous studies (Cui et al., 2025; Tan et al., 2020). Each slice was observed and photographed using microscope lenses at 1.25X and 20X magnification, respectively (Olympus B51167 Japan).

### 2.7 Cardiac Tissue RNA Extraction and Transcriptomics

Total RNA was extracted from cardiac tissue of 8-week-old male SHR fed a standard diet for 4 weeks (SHR group), 8-week-old adult SHR fed *red yeast rice* diet for 4 weeks (SHRD group), and 3-week-old SHR fed *red yeast rice* diet for 9 weeks (SHR-D group) using TRIzol reagent (ThermoFisher Scientific, USA) (n=4 per group). Library preparation and RNA sequencing were performed by GDIOD (Guangzhou, China).

### 2.8 Network pharmacology analysis and molecular docking analysis

Network pharmacology analysis and molecular docking analysis were performed as our previous study (Gilley et al., 2009; Nguyen et al., 2020; Tan et al., 2023).

### 2.10 Edu Analysis

H9c2 cells were procured from Shanghai Fuheng Biotechnology Co., Ltd. (Catalogue No. FH1004, China) and cultured in DMEM supplemented with 10% fetal bovine serum (FBS) in a 37°C, 5% CO₂ incubator. H9c2 cells were seeded into 12-well plates and cultured for 24 hours. They were then treated with Angiotensin II (Ang II, 200 nM) and low, medium, or high concentrations of MKA (5 μM, 50 μM, 500 μM) for 24 hours. Subsequently, Edu analysis was performed using EdU Apollo 567 In Vitro Kit (Absin (Shanghai) Biotechnology Co., Ltd., abs50050-200T) according to the manufacturer’s instructions. Images were observed and captured under a Leica fluorescence microscope. EdU-positive cells were quantified from images using Image J software.

### 2.11 TUNEL Analysis

Cells were fixed with 4% paraformaldehyde, permeabilized with 0.3% Triton X-100/PBS (VICMED) for 5 minutes, and processed using the One-step TUNEL In Situ Apoptosis Kit (Red, Elab Fluor® 594) (Elabscience Co., Ltd., Wuhan, E-CK-A322) according to the manufacturer’s instructions. Images were observed and captured under a Leica fluorescence microscope. Obtained images were analyzed by ImageJ.

### 2.12 Western blotting analysis

Total protein was extracted from H9c2 and rat heart tissues using RIPA lysis buffer containing PMSF and phosphatase inhibitors. Proteins (20–50 μg/well) were electrophoresed on a 10% sodium dodecyl sulphate (SDS) polyacrylamide gel and transferred to polyvinylidene fluoride (PVDF) membrane as described in our previous studies (Tan et al., 2020; Tan et al., 2023). Protein bands were visualized with ECL reagent (Vazyme Biotech, Nanjing, China) and analyzed by ImageJ Software.

### 2.13 Statistical Analysis

Image results obtained from experiments were analyzed and processed using ImageJ software. Statistical analysis was performed using GraphPad Prism software (version 10.0; GraphPad, USA). Data are presented as mean ± standard error (Mean ± SEM). Comparisons between two groups were performed using t-tests (and non-parametric tests), while comparisons among multiple groups were conducted using one-way analysis of variance (ANOVA) followed by Tukey’s multiple comparison test. Statistical significance was set at *p* < 0.05. More details are available in the Online Supplement.

## 3 Results

### 3.1 Manifestations of hypertension and cardiac hypertrophy in SHR

Following standard housing for 8, 12, and 16 weeks, systolic pressures and mean arterial pressures (MAP) were markedly elevated in SHR (**Figure 1A,B**). SHR exhibited significantly higher heart-to-body weight ratios and lung-to-body weight ratios compared to Control (**Figure 1C,D**). HE and Masson staining revealed markedly larger myocardial cell areas and fibrosis areas in SHR (**Figure 1E-G**). Echocardiography results demonstrated a time-dependent reduction in both the ejection fraction and shortening velocity of the SHR heart (**Figure 1H-J**). These findings indicate that 8-weeks SHR have exhibited significant hypertension, with myocardial hypertrophy and fibrosis progressively worsening over time.

**Figure 1.**
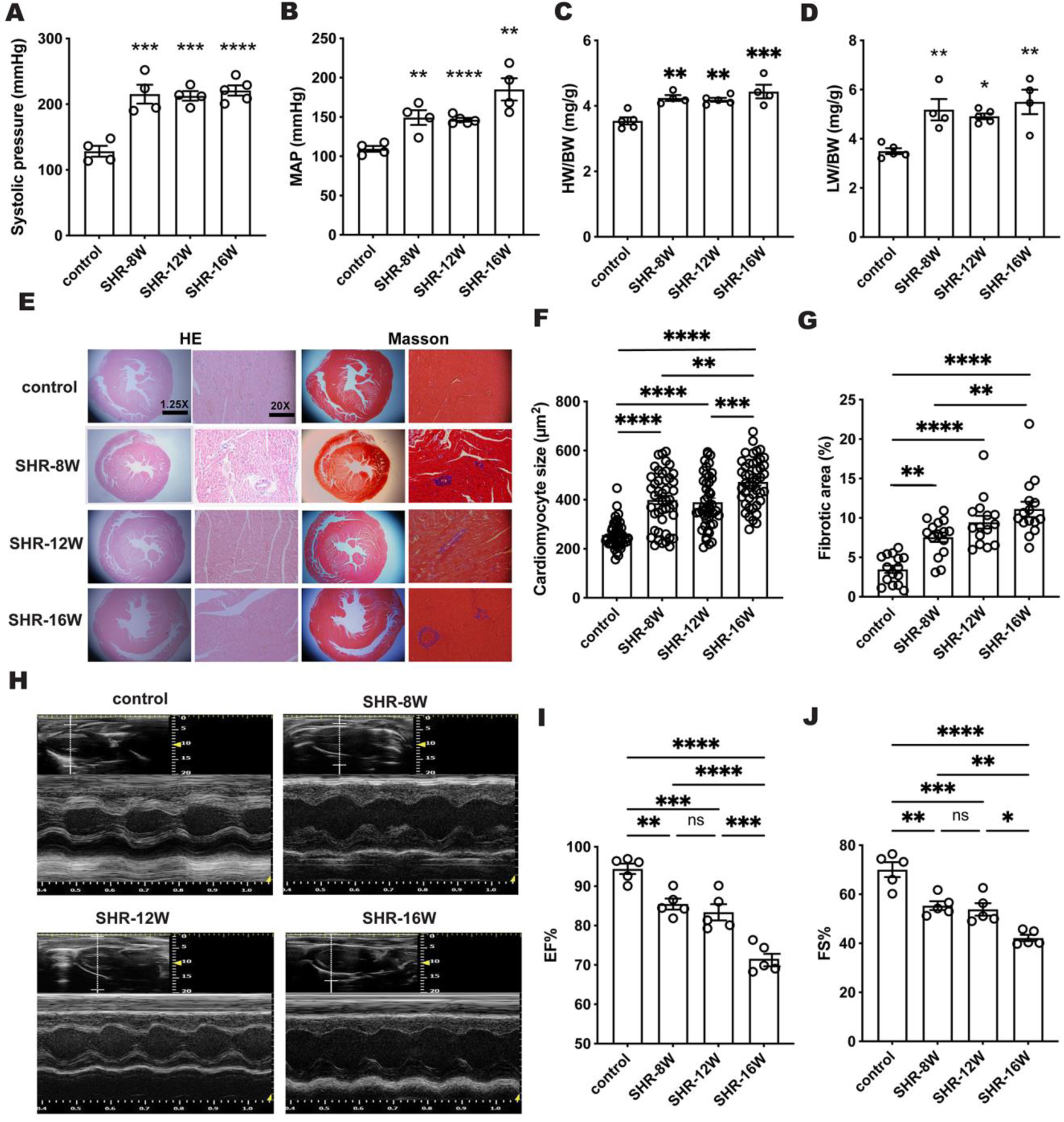
Cardiac structure and function in SHR. Systolic pressure (**A**) and mean arterial pressure (MAP, **B**) in 16-week Control rats versus 8-, 12-, and 16-week SHR rats. Heart-to-body weight ratio (**C**) and lung-to-body weight ratio (**D**) in 16-week Control rats versus 8-, 12-, and 16-week SHR rats. HE and Masson’s staining (**E**), myocardial cell cross-sectional area (**F**), and fibrotic area (**G**) in 16-week Control versus 8-, 12-, and 16-week SHR. Representative cardiac ultrasound (**H**), ejection fraction (EF%, **I**), and fractional shortening (FS%, **J**) in 16-week Control versus 8-, 12-, and 16-week SHR. Data are presented as mean ± SE, comparisons between two groups were performed using t-tests (and non-parametric tests), while comparisons among multiple groups were conducted using one-way analysis of variance (ANOVA) followed by Tukey’s multiple comparison test. (*p<0.05, **p<0.01, ***p<0.001, ****p<0.0001. n=3-5).

### 3.2 *Red yeast rice* significantly lowers blood pressure and alleviates cardiac remodeling in SHR

Following an 8-week period of feeding 8-week-old SHR chow diets (control) or chow diets containing 0.1%, 0.3%, and 0.6% red yeast rice, systolic pressures and MAP decreased markedly in 0.6% *red yeast rice* group compared with control group (**Figure 2A,B**), the heart-to-body weight ratio and lung-to-body weight ratio were also significantly reduced (**Figure 2C, D**). HE and Masson staining revealed that 0.6% *red yeast rice* inhibited cardiomyocytes hypertrophy and reduced fibrotic area (**Figure 2E-G**). Echocardiography revealed that the 0.6% *red yeast rice* treatment group exhibited significantly increased EF% and FS% (**Figure 2H-J**). Collectively, the cardioprotective effects of *red yeast rice* on SHR exhibited a concentration-dependent pattern, with the 0.6% concentration demonstrating optimal efficacy.

**Figure 2.**
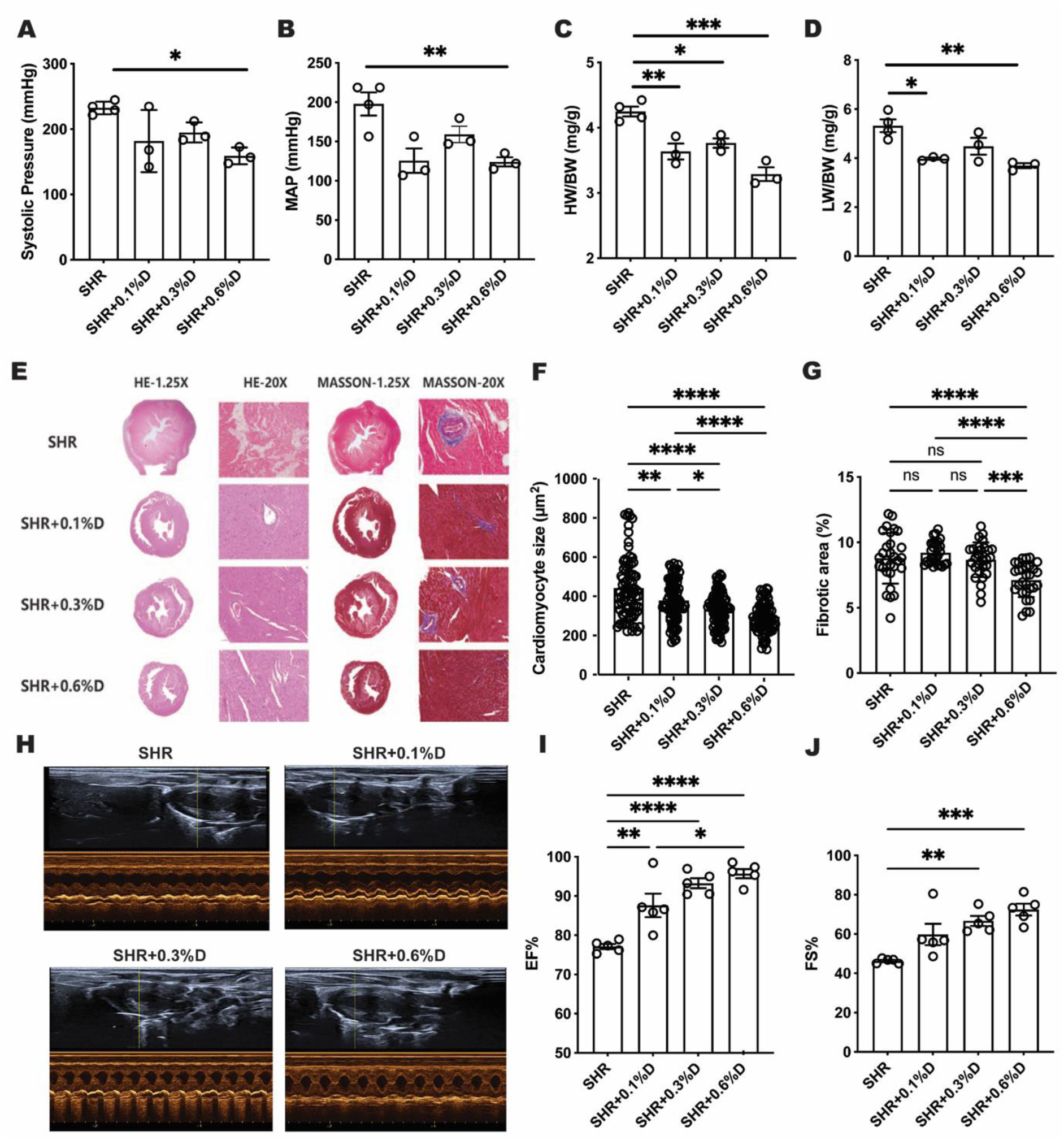
Cardiac structure and function in SHR fed red yeast rice diets at different concentrations. Systolic pressure (**A**), mean arterial pressure (MAP, **B**), Heart-to-body weight ratio (**C**), and lung-to-body weight ratio (**D**) in SHR fed chow diet (SHR) versus SHR fed 0.1%, 0.3%, and 0.6% red yeast rice diets (SHR+0.1%D, SHR+0.3%D, SHR+0.6%D). HE and Masson’s staining (**E**), myocardial cell cross-sectional area (**F**), and fibrotic area (**G**). Representative cardiac ultrasound (**H**), ejection fraction (EF%, **I**), and fractional shortening (FS%, **J**) in SHR, SHR+0.1%D, SHR+0.3%D, and SHR+0.6%D. Data are presented as mean ± SE, comparisons between two groups were performed using t-tests (and non-parametric tests), while comparisons among multiple groups were conducted using one-way analysis of variance (ANOVA) followed by Tukey’s multiple comparison test. (*p<0.05, **p<0.01, ***p<0.001, n=3-5).

Additionally, the therapeutic effects of 0.6% *red yeast rice* after 4-week versus 8-week treatment were compared. Results indicated no change in systolic pressure or MAP after 4 weeks, whereas they were markedly reduced after 8 weeks (***Figure S2A,B***). However, the cardiac weight-to-body weight ratio had already decreased significantly after 4 weeks of treatment (***Figure S2C,D***). HE and Masson staining revealed that cardiomyocytes hypertrophy and fibrosis were inhibited after 8 weeks of treatment (***Figure 2E-G***). This suggests that *red yeast rice* alleviates cardiac remodeling prior to blood pressure reduction. Consequently, we hypothesize that *red yeast rice* may directly target cardiac cells to improve myocardial remodeling.

### 3.3 Effects of *Red Yeast Rice* on blood pressure and cardiac remodeling in male and female SHR

It is well established that cardiovascular diseases exhibit sex differences. To investigate whether *red yeast rice* exerts distinct effects in male and female SHR, we selected age-matched male and female rats as subjects and fed them a 0.6% *red yeast rice* diet for 8 weeks. Results demonstrated significant reductions in systolic pressure and MAP in both female and male red yeast rice-treated groups (**Figure. 3A,B**). Heart-to-body weight ratio and lung-to-body weight ratio decreased (**Figure 3C,D**), whilst cardiac hypertrophy and fibrosis were suppressed (**Figure 3E-G**). No gender-specific differences in *red yeast rice* treatment efficacy were observed.

**Figure 3.**
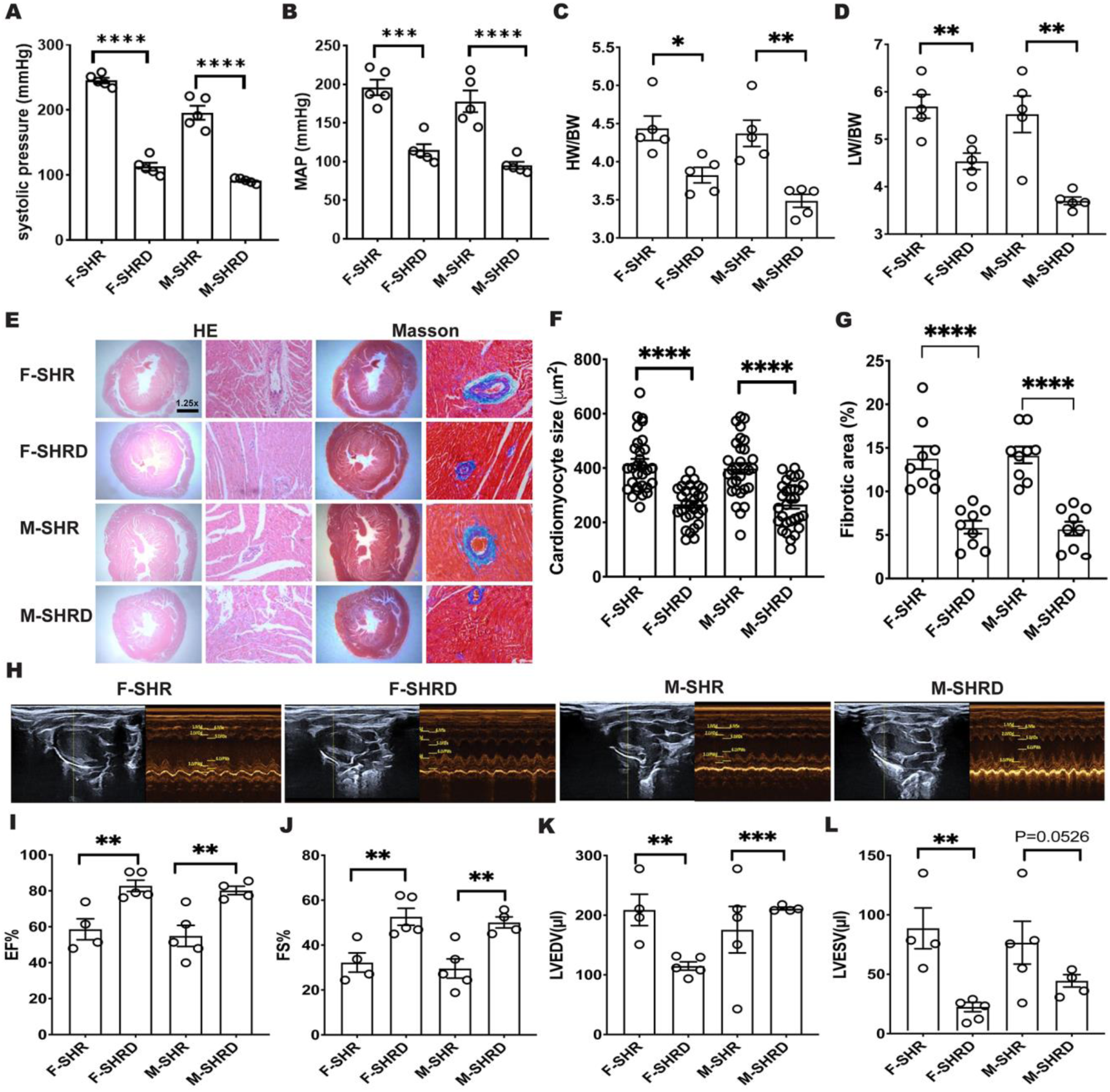
Cardiac structure and function in SHR fed red yeast rice diets at different gender. Systolic pressure (**A**), mean arterial pressure (MAP, **B**), Heart-to-body weight ratio (**C**), and lung-to-body weight ratio (**D**) in female SHR rats fed standard diet (F-SHR), female SHR rats fed red yeast rice diet (F-SHRD), male SHR rats fed standard diet (M-SHR), male SHR rats fed red yeast rice diet (M-SHRD). HE and Masson’s staining (**E**), myocardial cell cross-sectional area (**F**), and fibrotic area (**G**). Representative cardiac ultrasound (**H**), ejection fraction (EF%, **I**), fractional shortening (FS%, **J**), left ventricular end-diastolic volume (LVEDV, **K**), and left ventricular end-diastolic volume (LVESV, **L**) in F-SHR, F-SHRD, M-SHR, and M-SHRD. Data are presented as mean ± SE (*p<0.05, **p<0.01, ***p<0.001, n=3-5).

Echocardiographic findings revealed significantly increased EF% and FS% in both female and male red yeast rice-treated groups (**Figure 3H-J**), alongside markedly reduced LVEDV and LVSDV (**Figure 3K,L**). The therapeutic effects on cardiac function showed no gender-specific differences.

To evaluate potential sex-specific toxicological differences in vital organs, histological staining analysis was performed on lung, liver, and kidney tissues. Results demonstrated that *red yeast rice* significantly ameliorated fibrosis and inflammation in lung, liver, and kidney tissues of both male and female SHR, with no observed sex differences (***Figure S3***). No other toxicological alterations were detected.

### 3.4 Effects of *Red Yeast Rice* on blood pressure and cardiac remodeling in progeny SHR

Extensive research indicates that the health status, diet, and medication of progenitors influence offspring health. However, whether administering *red yeast rice* to progenitor SHR may cause adverse effects in their progeny, or whether it confers benefits to offspring blood pressure and cardiac function, warrants investigation.

Progenitors were divided into red yeast rice-treated and standard diet groups. Their offspring were categorized as: neither progenitors nor offspring treated (SHR); progenitors untreated, offspring treated (SHR+D); progenitors treated, offspring untreated (SHRD+N); and both progenitors and offspring treated (SHRD+D). At eight weeks of age, offspring underwent hemodynamic testing and tissue section staining. The results revealed significantly reduced systolic pressure and MAP across all treatment groups compared to SHR, with SHRD+D and SHRD+N demonstrating greater reductions than SHR+D (**Figure 4A-B**). The SHRD+D group exhibited a marked decrease in the heart-to-body weight ratio (**Figure 4C-D**). HE and Masson staining revealed the most pronounced reduction in myocardial cell area and fibrotic area within the SHRD+D group (**Figure 4E-G**).

**Figure 4.**
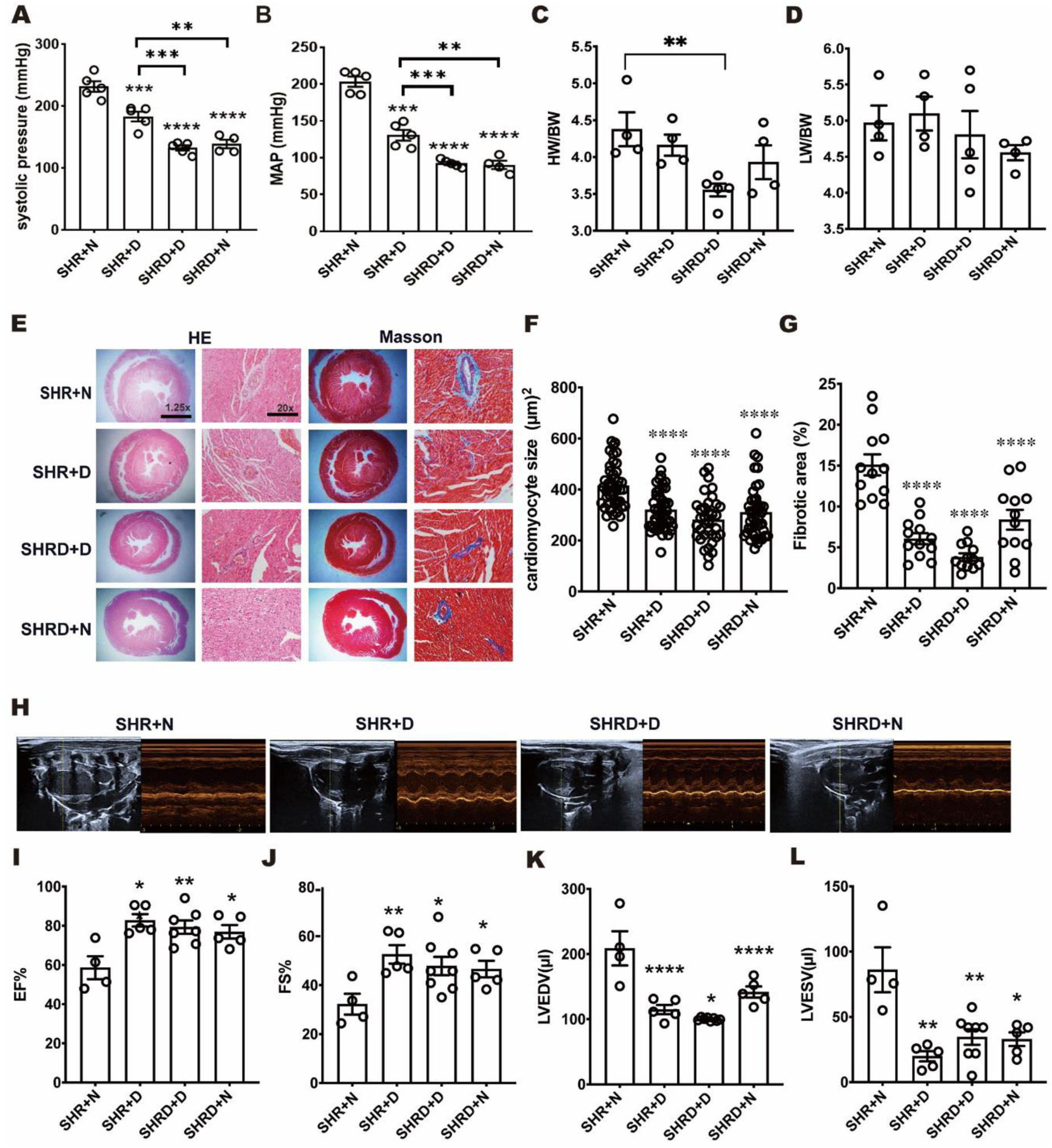
Cardiac structure and function in SHR fed red yeast rice diets at different offspring. Systolic pressure (**A**), mean arterial pressure (MAP, **B**), Heart-to-body weight ratio (**C**), and lung-to-body weight ratio (**D**) in parent SHR fed chow diet produced offspring also fed chow diet (SHR+N); parent SHR fed chow diet produced offspring fed red yeast rice diet (SHR+D); parent SHR fed red yeast rice diet produced offspring also fed red yeast rice diet (SHRD+D); parental SHR fed red yeast rice diet produced offspring SHR fed chow diet (SHRD+N). HE and Masson’s staining (**E**), myocardial cell cross-sectional area (**F**), and fibrotic area (**G**). Representative cardiac ultrasound (**H**), ejection fraction (EF%, **I**), fractional shortening (FS%, **J**), left ventricular end-diastolic volume (LVEDV, **K**), and left ventricular end-diastolic volume (LVESV, **L**) in SHR+N, SHR+D, SHRD+D, and SHRD+N. Data are presented as mean ± SE, comparisons between two groups were performed using t-tests (and non-parametric tests), while comparisons among multiple groups were conducted using one-way analysis of variance (ANOVA) followed by Tukey’s multiple comparison test (*p<0.05, **p<0.01, ***p<0.001, n=3-5).

Echocardiography demonstrated significant increased EF% and FS% alongside reduction in LVEDV in the SHRD+D group compared to other groups (**Figure 4H-J**). These findings suggest parental drug administration confers beneficial effects on offspring blood pressure and cardiac function, with sustained treatment yielding superior outcomes.

Furthermore, to assess potential adverse effects on other vital organs in offspring, histological staining analyses were conducted on lung, liver, and kidney tissues. Results demonstrated that *red yeast rice* significantly ameliorated fibrosis and inflammation in the lungs, liver, and kidneys of offspring SHR, with the SHRD+D group exhibiting the most pronounced effects (***Figure S4***). No other toxicological alterations were observed.

### 3.5 Transcriptomics combined with network pharmacology to identify the molecular mechanism

To further investigate the molecular mechanism by which *red yeast rice* ameliorates myocardial hypertrophy, we employed transcriptomics combined with network pharmacology to screen the targets and signaling pathways of MKA (3.5mg/g), the primary active component of red yeast rice. The two-dimensional molecular structure of MKA is depicted in **Fig. 5A**. Following target prediction for MKA using the Swiss-Target Prediction database, 100 potential targets were identified. Extensive research indicates that MKA can significantly reduce LDL-C. However, there are no reports indicating whether MKA directly targets cardiomyocytes to improve hypertensive cardiac hypertrophy.

**Figure 5.**
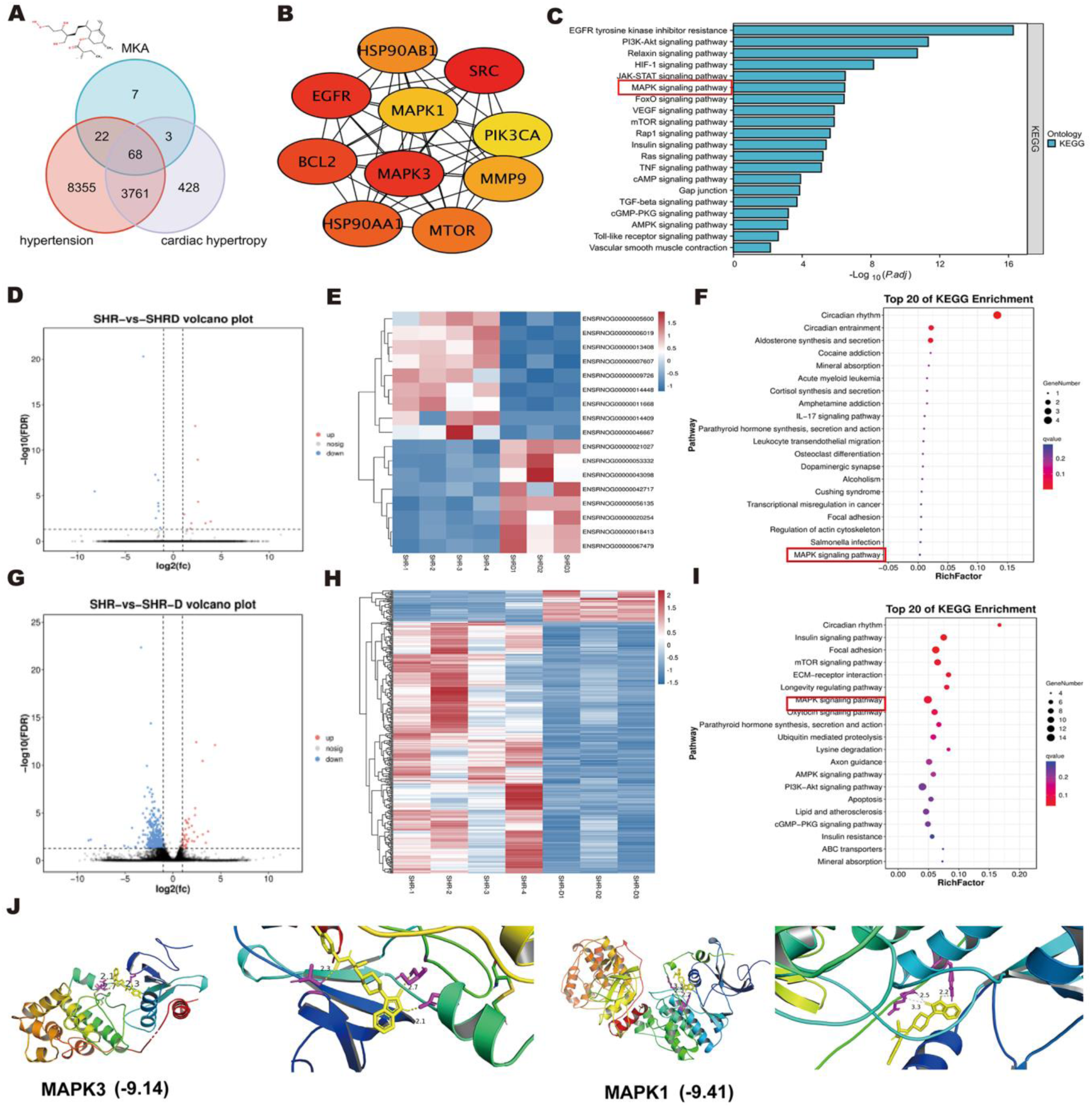
Network pharmacology combined with transcriptomics identifies the molecular mechanism of MKA action in red yeast rice. **A.** Venn diagram of MKA targets and genes associated with hypertension and myocardial hypertrophy. **B.** Top 10 central genes among 71 common target genes for MKA and myocardial hypertrophy. **C.** KEGG pathway analysis results for MKA-targeted cardiac hypertrophy genes. **D.** Volcano plot of differentially expressed genes comparing 8-week-old male SHR fed a standard diet for 4 weeks (SHR group), 8-week-old adult SHR fed red yeast rice diet for 4 weeks (SHRD group). **E.** Heatmap of differentially expressed genes comparing SHR and SHRD groups. **F.** KEGG pathway analysis results for differentially expressed genes comparing SHR and SHRD groups. **G.** Volcano plot of differentially expressed genes comparing SHR and 3-week-old SHR fed red yeast rice diet for 9 weeks (SHR-D group). **H.** Heatmap of differentially expressed genes comparing SHR and SHR-D groups. **I.** KEGG pathway analysis results for differentially expressed genes comparing SHR and SHR-D groups.

Using the keyword “hypertension”, 12,116 hypertension-related targets were obtained from the GeneCard database. A total of 635 target genes with a score ≥8 were selected. The Venn diagram revealed 90 common target genes between MKA and hypertension (**Fig. 5A**). Using the keyword “myocardial hypertrophy”, 4,189 targets associated with myocardial hypertrophy were obtained. A total of 1,719 target genes with a score ≥8 were selected. The Venn diagram revealed 71 common target genes between MKA and myocardial hypertrophy (**Figure 5A**). Cytoscape analysis identified the top 10 potential targets (**Figure 5B**): *EGFR*, *MAPK3*, *MMP9*, *SRC*, *MAPK1*, *BCL2*, *HSP90AA1*, *HSP90AB1*, *PIK3CA*, and *MTOR*. KEGG pathway analysis revealed key signaling pathways including EGFR, PI3K-Akt, MAPK, and so on (**Figure 5C**). Volcano plots and heatmaps derived from transcriptomic analysis reveal that differentially expressed genes upregulated or downregulated in the SHR group compared to the SHR-D/SHRD group (**Figure 5D,E,G,H**). Furthermore, the number of differentially expressed genes markedly increased when comparing the SHR group to the SHR-D group. It is evident that as the duration of drug administration increased, the number of differentially expressed genes in rats at the same gestational age rose, and the magnitude of transcriptional changes became more pronounced.

Subsequently, KEGG analysis of the differentially expressed genes yielded the top 20 signaling pathways (**Figure 5F, I**). Integrating data from both the SHR and SHRD groups and the SHR and SHR-D groups revealed that the MAPK pathway was consistently implicated in both sets of analyses.

Therefore, we hypothesize that MKA inhibits hypertension and cardiac hypertrophy by targeting MAPK1 (ERK2) and MAPK3 (ERK1). Binding energy is considered robust at ≤-5.0 kcal/mol and exceptionally strong at ≤-7.0 kcal/mol. Molecular docking results demonstrate that MKA exhibits strong binding affinity with both MAPK1 (ERK2) and MAPK3 (ERK1) proteins (affinity values of -9.41 kcal/mol and -9.14 kcal/mol, **Figure 5J**). These findings indicate that MKA functions by forming stable complexes with common target genes of MAPK1 (ERK2) and MAPK3 (ERK1).

### 3.6 *Red Yeast Rice* targets ERK1/2 to downregulate c-Fos and improve cardiac hypertrophy markers in SHR

The effects of *red yeast rice* treatment on SHR were investigated using Western blot analysis. *Red yeast rice* (0.6%) reduced expression of myocardial fibrosis markers Col1α and FN1, cardiac hypertrophy protein β-MHC, and apoptotic protein Caspase 3 (*p < 0.05, **p < 0.01, ***p < 0.001, ****p < 0.0001, **Figure 6A-E**). To verify that MKA can act on MAPK1 (ERK2) and MAPK3 (ERK1) to inhibit hypertension and cardiac hypertrophy, we checked the level of p-ERK1/2 in the cardiac tissue between SHR group and SHRD group. Western blot results demonstrated that after eight weeks of *red yeast rice* diet administration, the expression levels of p-ERK1/2 in cardiac tissue were markedly suppressed (**Figure 6F,G**). Transcriptomic analysis further revealed that expression of the transcription factor c-Fos was significantly downregulated in the treatment group (**Figure 6H**). Western blot results confirmed that *red yeast rice* significantly downregulated the expression of c-Fos (**Figure 6G**).

**Figure 6.**
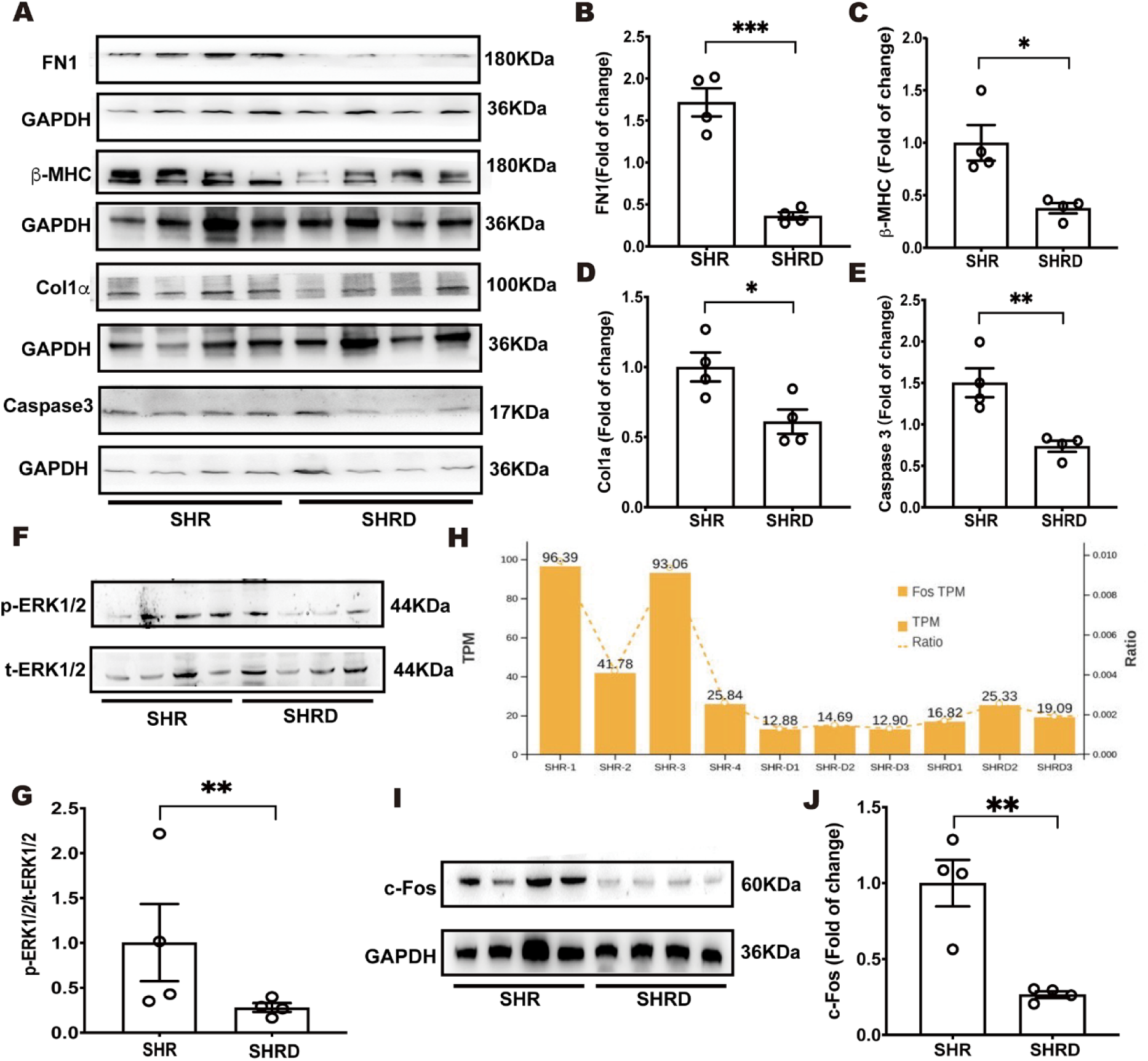
Western blot results for cardiac tissue proteins in SHR following red yeast rice treatment. **A-E.** Representative Western blot bands and relative protein expression for FN1, β-MHC, COL1α, Caspase-3 and GADPH; **F,G.** Representative Western blot bands and relative protein expression for p-ERK and t-ERK, **H.** Transcriptomic results of transcription factor c-Fos. **I,J.** Representative Western blot bands and relative protein expression for c-Fos, Data are presented as mean ± SE, comparisons between two groups were performed using t-tests (and non-parametric tests) (*p < 0.05, **p < 0.01, ***p < 0.001, n=4).

Previous studies have demonstrated that nuclear translocation of phosphorylated ERK1/2 promotes the transcriptional activity c-Fos (Ren et al., 2023). We therefore hypothesize that red yeast rice, through its primary active component MKA, targets ERK1/2 by inhibiting its phosphorylation, suppresses c-Fos expression, ultimately inhibiting downstream hypertrophic gene expression and thereby mitigating cardiac hypertrophy in SHR.

### 3.7 MKA reverses Ang II-induced cardiomyocyte injury by inhibiting ERK1/2/c-Fos

Ang II treatment of H9c2 cardiomyocytes is widely employed to simulate an in vitro model of hypertensive cardiac hypertrophy (Tham et al., 2015). Following 24-hour treatment of H9c2 cardiomyocytes with 200 nM Angiotensin II, EDU cell staining analysis revealed that Ang II significantly suppressed the viability of H9c2 cardiomyocytes (**Fig. 7A**). Concurrent administration of 1 μM, 5 μM, and 10 μM MKA for 24 hours revealed that the 5 μM group exhibited significantly reversed cell viability, indistinguishable from the control group (**Fig. 7A,B**). TUNEL staining analysis revealed that Ang II significantly induced apoptosis in H9c2 cardiomyocytes, while 5 μM MKA markedly suppressed Ang II-induced apoptosis (**Figure 7A,C**).

**Figure 7.**
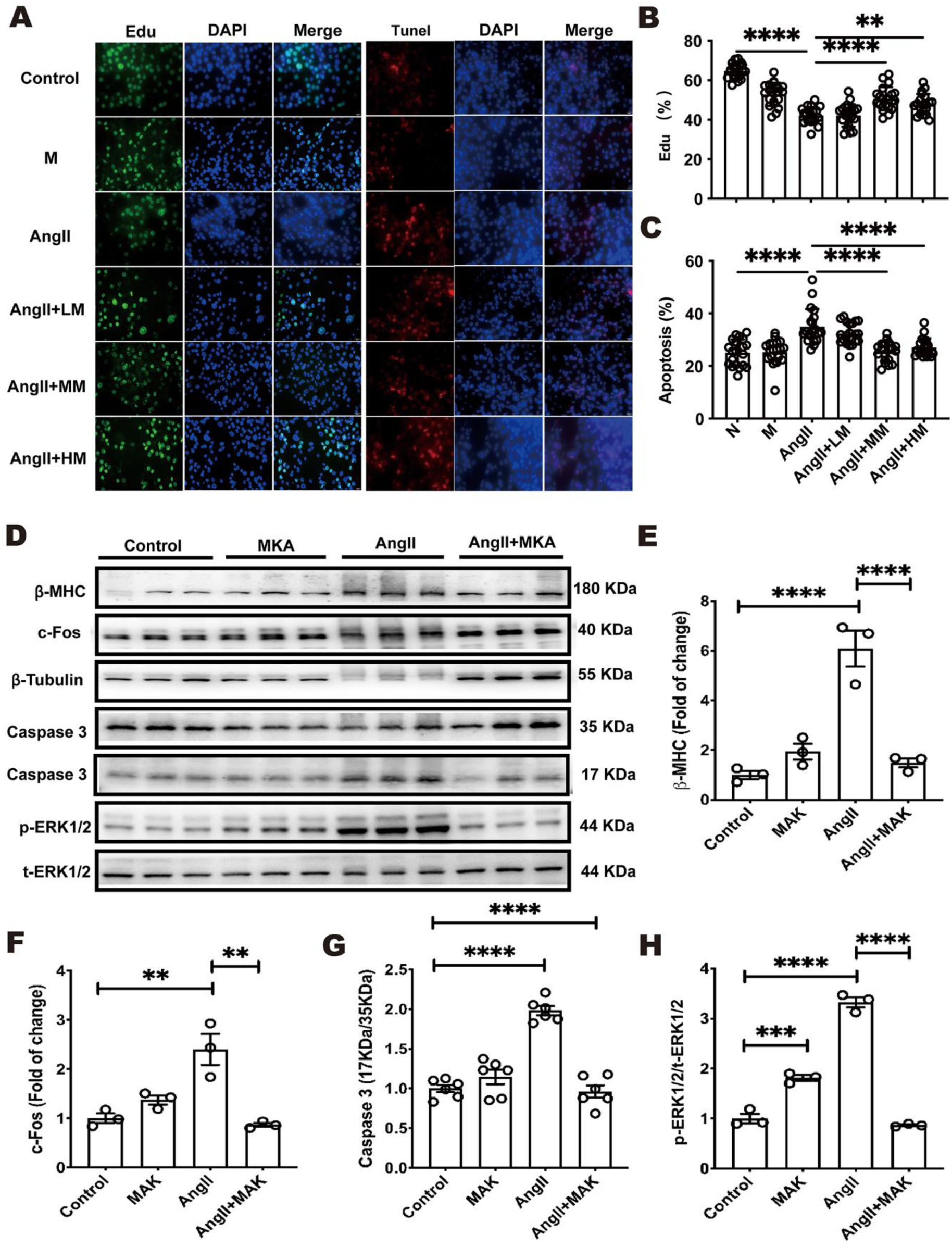
Effects of MKA on Angiotensin II-induced cardiomyocyte injury and hypertrophy. Representative images of EDU and TUNEL staining (**A**), EDU positive percent (**B**) and apoptosis percent (**C**) in H9c2 cardiomyocytes treated with different concentrations of MKA (1, 5, 10 μM) and 200 nM Ang II for 24 hours. **D-H.** Representative Western blots and relative protein expression for β-MHC, p-ERK, t-ERK, c-Fos, and Caspase-3. Data are presented as mean ± SE, comparisons among multiple groups were conducted using one-way analysis of variance (ANOVA) followed by Tukey’s multiple comparison test (n = 4 per group; *p < 0.05, **p < 0.01, ***p < 0.001, ****p < 0.0001). LM: 1 μM MKA; MM: 5 μM MKA; HM: 10 μM MKA;

Western blot analysis validated MKA’s effects on hypertrophy and apoptosis-related proteins in AngII-treated H9c2 cardiomyocytes. Ang II significantly upregulated β-MHC and caspase-3 expression (**Figure 7D-H**), which was reversed following 5 μM MKA treatment. Furthermore, Ang II markedly increased p-ERK1/2 and c-Fos expression, which was completely reversed by 5 μM MKA treatment (**Figure 7F,H**). We therefore hypothesize that MKA downregulates c-fos expression by targeting and inhibiting phospho-ERK1/2, leading to significant suppression of β-MHC and caspase-3 expression and ultimately inhibiting myocardial hypertrophy.

## 4. Discussion

Left ventricular hypertrophy is the most common clinical manifestation of hypertension, resulting from cardiac cellular remodeling and enlargement. It constitutes a potent risk factor for cardiovascular events and all-cause mortality, particularly arrhythmias and sudden cardiac death (Okin et al., 2013). Compared with individuals without left ventricular hypertrophy, the presence of hypertrophy increases the risk of ventricular arrhythmias by 5.5% and that of sudden cardiac death by 1.2%, ventricular tachycardia/ventricular fibrillation by 2.8-fold (Tin et al., 2002), and atrial fibrillation by 40–50% (Sirtori, 2014). Statins are commonly used adjunctive medications for hypertension management. Beyond cholesterol reduction, statins lower blood pressure through multiple mechanisms, including improved vascular endothelial function, reduced vascular inflammation, and decreased sympathetic nervous system activity. Additionally, statins may improve insulin resistance, thereby contributing to blood pressure reduction (Alonso et al., 2019). Our research also found that red yeast rice containing MKA can alleviate hypertensive cardiac hypertrophy.

Given the marked differences in hypertension between genders, particularly regarding age of onset and medication use, gender represents another crucial determinant requiring assessment. A large meta-analysis of 46 population-based studies across 22 countries, involving 123,143 men and 164,858 women aged 20–59 years, indicated that hypertensive women were 1.33 times more likely than men to receive drug therapy, with a higher propensity for diuretic use, whereas men more frequently utilized beta-blockers, ACE inhibitors, and calcium channel blockers (Busjahn et al., 1996). Among patients receiving monotherapy for hypertension, women were less likely than men to be prescribed beta-blockers, calcium channel blockers, or ACE inhibitors (Os et al., 1994). One possible explanation for these findings may be gender differences in side effects. Women are more prone to coughing when treated with ACE inhibitors compared to men (Agrawal and Wenger, 2020). The results of this study indicate no significant gender differences in the therapeutic efficacy of red yeast rice, thereby providing a research basis for the potential administration of identical medications across different genders.

Research indicates that parental health profoundly and persistently influences numerous aspects of offspring development in utero and subsequent physiological and metabolic outcomes (Bramham et al., 2014). Among women with chronic hypertension, fetal growth restriction is observed in 10-20% of pregnancies (Kattah and Garovic, 2013), higher rates of preterm birth before 37 weeks gestation, lower birth weight, neonatal unit admissions, and perinatal mortality (McCowan et al., 1996). Two primary oral antihypertensive agents are available for pregnant women: labetalol and nifedipine (Tomar et al., 2024). However, neither of these drugs can directly improve cardiac hypertrophy. Our study indicated that maternal consumption of *red yeast rice* during pregnancy does not impair fetal growth and development, while conferring cardiac benefits to the offspring postnatally. Concurrently, research demonstrates that beyond maternal health influencing the fetus, the paternal dietary patterns and health status prior to conception exert significant effects on offspring metabolism (Aiken et al., 2016). For instance, paternal overweight at conception doubles offspring obesity risk and impairs metabolic health (Xu et al., 2021). Acute high-fat diet feeding or genetically induced mitochondrial dysfunction in male mice resulted in impaired glucose homeostasis in male offspring (Aiken et al., 2016). This suggests that the health of hypertensive fathers also affects progeny. Consequently, this study concurrently administered *red yeast rice* intervention to hypertensive male rats to observe effects on offspring. Results indicate that pre-pregnancy *red yeast rice* intervention in both parental rats confers cardiac benefits. Continued postnatal administration to offspring further enhances these effects, providing experimental evidence for the safety of *red yeast rice* consumption during pre-pregnancy and lactation in hypertensive parents and its associated benefits for offspring.

Increasing evidence indicates that the MAPK/ERK1/2 signaling pathway is one of the key pathways contributing to cardiac hypertrophy (Dash et al., 2001), and its targeted inhibition can ameliorate hypertrophy. Song et al. (2024) found that artemisinin alleviated isoproterenol induced cardiac hypertrophy through inhibiting the ERK1/2 signaling pathways. In clinical trials, the MAPK pathway has seen preliminary therapeutic application. Flesch et al. (2001) employed mechanical unloading of the heart via a left ventricular assist device, demonstrating that mechanical unloading of failing hearts improves cardiac function through differential regulation of the MAPK pathway. Babu et al. (2000) utilized deoxyepinephrine-induced hypertrophic cardiomyocytes to demonstrate that phosphorylated ERK1/2 binds to c-Fos, ultimately leading to excessive cardiomyocyte proliferation. These findings collectively highlight the substantial potential of the ERK1/2/c-Fos pathway in ameliorating myocardial hypertrophy. Meantime, our study clarified that *red yeast rice* derived MKA attenuates cardiac hypertrophy in SHR by targeting the ERK1/2/c-Fos signaling axis. This mechanism transcends the limitations of traditional *red yeast rice* research, which has primarily focused on lipid-lowering effects, offering fresh perspectives on its cardiovascular protective functions.

This study remains subject to several limitations. Firstly, mechanism validation was confined to rat models and H9c2 cells; the conservation of the ERK1/2/c-Fos pathway requires verification in primary human cardiomyocytes or organoid models. Secondly, the mechanism underpinning the transgenerational protective effect remains unclear, necessitating investigation into whether epigenetic regulation (such as DNA methylation or non-coding RNAs) is involved. Thirdly, long-term clinical trials are required to evaluate red yeast rice’s efficacy in reversing human left ventricular hypertrophy and its impact on cardiovascular event endpoints. Fourthly, as a traditional Chinese medicine, our study focused solely on the action and molecular mechanisms of MKA, the primary component of red yeast rice, without investigating the effects of other constituents. Fifthly, the long-term safety profile of this herbal medicine warrants further investigation. Sixthly, future research should examine the combined effects of *red yeast rice* with other cardiovascular medications to optimize therapeutic regimens.

## Conclusion

This study confirms that MKA derived from *red yeast rice* improves hypertensive cardiac hypertrophy by blocking c-Fos-mediated hypertrophic gene expression through targeted binding to ERK1/2 and inhibiting its phosphorylation (**Figure 8**). Its demonstrated therapeutic stability in animal models, multi-organ safety profile, and transgenerational protective effects provide substantial support for *red yeast rice* as a dietary intervention strategy for hypertensive cardiac hypertrophy.

**Figure 8.**
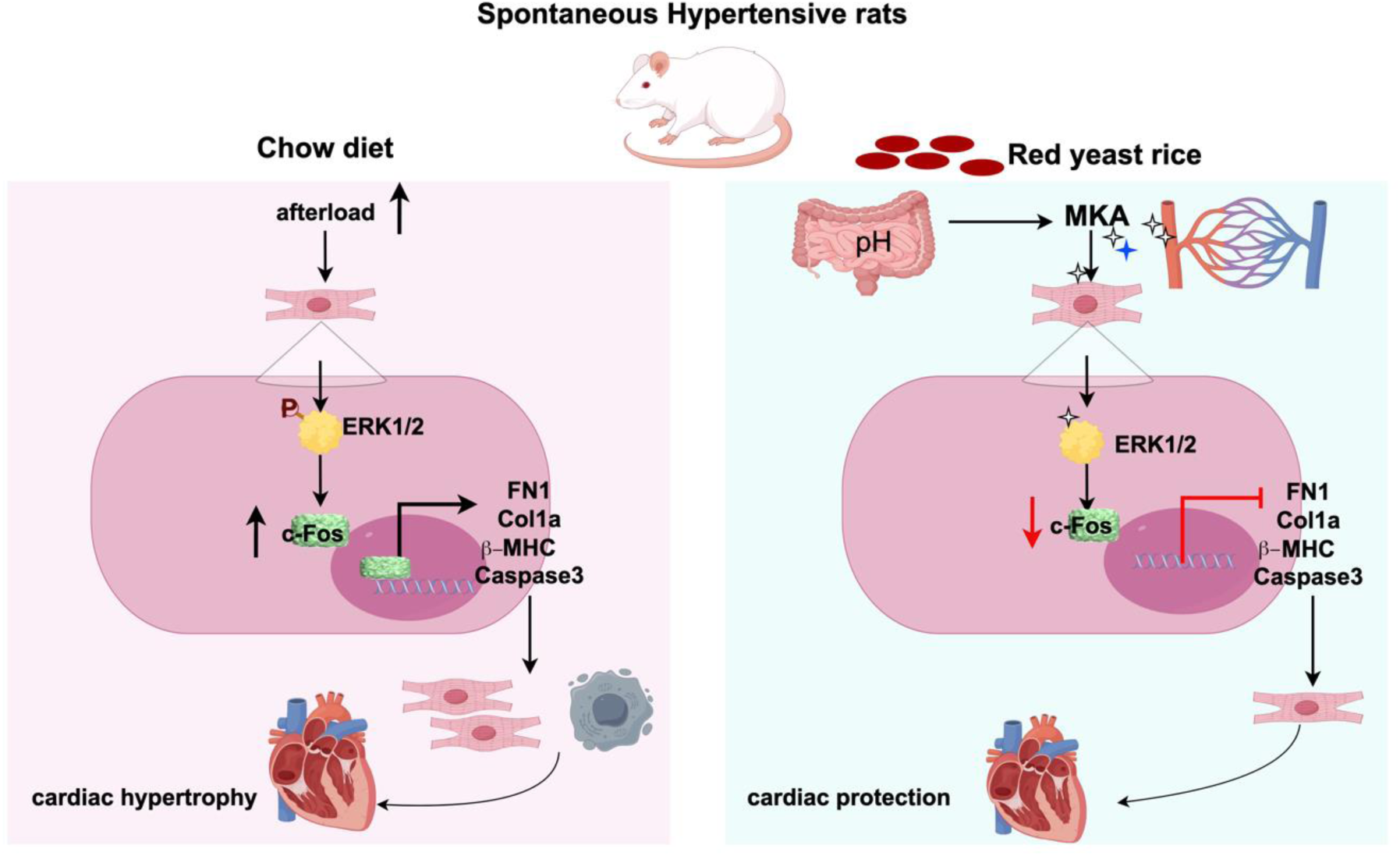
The mechanism of red yeast rice derived MKA on hypertension-induced cardiac hypertrophy.

## Abbreviation

ACE: Angiotensin converting enzyme
ACEIs: Agents acting on the renin-angiotensin system
Ang: II Angiotensin II
β-MHC: Beta-myosin heavy chain
BCL2: B-cell lymphoma-2
c-Fos: AP-1 transcription factor subunit
caspase 3: Cysteinyl aspartate specific proteinase 3
CCBs: Calcium channel blockers
Col1α: Collagen type 1 α
DMEM: Dulbecco’s modified Eagle’s medium
EGFR: Epidermal growth factor receptor
ERK1/2: Extracellular signal-regulated kinase1/2
FBS: Fetal bovine serum
FN1: Fibronectin 1
GAPDH: Glyceraldehyde 3-phosphate dehydrogenase
H&E: Hematoxylin & eosin
HRP: Horseradish peroxidase
LDL-C: Low-density lipoprotein cholesterol
LVEDV: Left ventricular end-diastolic volume
LVESV: Left ventricular end-systolic volume
LVH: Left ventricular hypertrophy
MAP: Mean arterial pressures
MAPK: Mitogen-activated protein kinases
MKA: Monacolin K β-hydroxy acid
MMP9: Matrix metalloprotease 9
MTOR: Mammalian target of rapamycin
non-HDL: Non-high-density lipoprotein
p-ERK: Phosphorylated extracellular signal-regulated kinase
PDB: Protein Data Bank
PI3K: Phosphatidylinositol-4,5-bisphosphate 3-kinase
PVDF: Polyvinylidene fluoride
SHR: Spontaneously hypertensive rats
t-ERK: Total extracellular signal-regulated kinase

## CRediT authorship contribution statement

**Rubin Tan:** Writing – original draft, Writing – review & editing, Software, Methodology, Investigation, Formal analysis, Data curation, Conceptualization, Supervision, Conceptualization, Project administration, Funding acquisition. **Dongqi Yang:** Writing – original draft, Software, Formal analysis, Data curation. **Kuntao Liu:** Writing – original draft, Software, Formal analysis, Data curation. **Jia Liu:** Formal analysis, Data curation, Writing – original draft. **Na Li:** Formal analysis, Data curation, Writing – original draft. **Jie Cui:** Data curation. **Xiaoqiu Tan:** Supervision, Funding acquisition, Writing – review & editing. **Qinghua Hu:** Supervision, Project administration, Funding acquisition, Writing – review & editing. **Chunxiang Zhang:** Supervision, Project administration, Funding acquisition, Writing – review & editing. All authors read and approved the final manuscript.

## Acknowledgments

This work was supported by grants from the National Natural Science Foundation of China (81700055, U23A20398, 82030007, 82270334 and 82470323), Noncommunicable Chronic Diseases-National Science and Technology Major Project (2024ZD0537707), the Science and Technology Department of Sichuan Province (2025ZNSFSC0052), and the Natural Science Foundation of Jiangsu Province (BK20160229), and Southwest Medical University Talent Launch Project (25110200005). The authors would like to express their gratitude to Kexue Li and Jingxia Kuai at Xuzhou Medical University for their assistance in our research.

## Data Availability Statement

The data that support the findings of this study are available from the corresponding author upon reasonable request.

